# Correlation of Microglial Activation with White Matter Changes in Dementia with Lewy Bodies

**DOI:** 10.1101/599241

**Authors:** Nicolas Nicastro, Elijah Mak, Guy B. Williams, Ajenthan Surendranathan, William Richard Bevan-Jones, Luca Passamonti, Patricia Vazquez Rodriguez, Li Su, Robert Arnold, Franklin I. Aigbirhio, James B. Rowe, John T. O’brien

## Abstract

Dementia with Lewy bodies (DLB) is the second-leading degenerative dementia after Alzheimer’s disase. Neuropathologically, it is characterized by alpha-synuclein protein deposition with variable degree of concurrent Alzheimer pathology. Neuroinflammation is increasingly recognized as a significant contributor of degeneration.

**Objective:** to examine the relationship between microglial activation as measured with [^11^C]-PK11195 brain PET and MR diffusion tensor imaging (DTI) in DLB.

**Methods:** nineteen clinically probable DLB and 20 similarly aged controls underwent structural MRI with T1-weighted and 3T DTI sequences. Eighteen DLB subjects also underwent [^11^C]-PK11195 PET imaging. Tract-Based Spatial Statistics (TBSS) were performed to compare DTI parameters in DLB relative to controls and identify associations of [^11^C]-PK11195 binding with white matter integrity.

**Results:** TBSS showed widespread changes in all DTI parameters in the DLB group compared to controls (Threshold Free Cluster Enhancement (TFCE) p < 0.05). [^11^C]-PK11195 binding in parietal cortices also correlated with widespread lower mean and radial diffusivity (TFCE p < 0.05).

**Conclusion:** Our study demonstrates that higher PK11195 binding is associated with a relative preservation of white matter, positioning neuroinflammation as a potential early marker in the DLB pathogenic cascade.

## INTRODUCTION

Dementia with Lewy bodies (DLB) is the second-leading degenerative dementia in older people^1^. It is characterized by alpha-synuclein protein deposition in the form of intraneuronal Lewy bodies as well as Lewy neurites, with variable degree of concurrent Alzheimer pathology^2, 3^. Neuroinflammation and, more specifically microglial activation, is increasingly recognized as a key etiopathogenic mechanism in DLB and can be assessed in vivo via the use of different PET tracers such as [^11^C]-PK11195 (PK11195)^4^. In DLB, increased microglial activation as measured with PK11195 has been observed in striatum, substantia nigra and widespread cortical regions^5, 6^. Intriguingly, we recently showed that subjects with a cognitively mild DLB had higher PK11195 binding than those with a moderate impairment in regions including frontal, temporal and parietal cortices, caudate nucleus and thalamus^6^. Recent studies have shown that microglial activation is an initial event in Alzheimer’s disease (AD), mirroring the neuropathological changes primarily involving temporoparietal regions^7, 8^. However and at variance with data from patients with clinical AD^9^, amyloid burden in DLB, measured with ^11^C-Pittsburgh Compound B (PiB), was not correlated with PK11195 binding^6^. Together, these observations suggest a key role for microglial activation in DLB, adding credence to the view that neuroinflammation may represent a potential therapeutic target in the field of dementia. Unlike molecules targeting protein accumulation which are currently at an experimental stage, anti-inflammatory drugs are already available and could be brought more quickly into clinical drug trials. However, this would first require a better understanding of the temporal ordering and clinical relevance of protein accumulation, neuroinflammation and structural brain changes. White matter abnormalities, especially reduced fractional anisotropy (FA) and increased mean and radial diffusivity (MD and RD, respectively), are recognised features in DLB, especially in the absence of the marked brain atrophy seen in AD. Several pathological mechanisms for such white matter changes have been described in dementia, including ischaemia, demyelination, amyoid angiopathy and inflammation.

In the present study, we aimed to characterize the relationship between microglial activation as measured with PK11195 PET and diffusion tensor imaging (DTI) impairment in a cross-sectional group of DLB subjects in which the amyloid status had been confirmed through PiB PET imaging^6^. As we recently observed that DLB subjects with a milder cognitive impairment had widespread higher cortical PK11195 uptake, especially in frontal and temporal lobes, we hypothesized that microglial activation in these regions correlates with a relative preservation of white matter microstructure.

## METHODS

### Participants

The present work is part of the Neuroimaging of Inflammation in MemoRy and Other Disorders (NIMROD) study^10^. All participants were aged over 50 years and had sufficient proficiency in English for cognitive testing. We included 19 participants with probable DLB according to both 2005 and 2017 consensus criteria^11, 12^ as described in ^6^. In addition, subjects had at least two years clinical follow-up to confirm clinical progression and no change of diagnosis. We also recruited 20 similarly aged healthy controls, with MMSE scores greater than 26, absence of regular memory complaints, and no unstable or significant medical illnesses. A detailed clinical and neuropsychological assessment was performed^10^.

Patients were identified from the Memory clinic at the Cambridge University Hospitals NHS Trust, other local memory clinics, and from the Dementias and Neurodegenerative Diseases Research Network (DeNDRoN) volunteer registers. Healthy controls were recruited via DeNDRoN as well as from spouses and partners of participants. Informed written consent was obtained in accordance with the Declaration of Helsinki. The study received a favourable opinion from the East of England Ethics Committee (Cambridge Central Research, Ref. 13/EE/0104).

### PET acquisition and MRI

Among 19 DLB subjects with available DTI imaging, 18 participants underwent PET imaging on a GE Advance or GE Discovery 960 scanner with the radioligand [^11^C]-PK11195 and also underwent multi-modal 3T MPRAGE and DTI imaging as described in detail below. In addition, 15 DLB subjects underwent [^11^C]-PiB PET imaging to quantify the density of fibrillar Aβ deposits for classification of Aβ status (positive when PiB cortical standardized uptake value ratio (SUVR) > 1.5)^13^. The [^11^C]-PK11195 PET image series were aligned across the frames to correct for head motion during data acquisition with SPM12. The realigned dynamic frames were co-registered to the T1-MPRAGE, which were processed with Freesurfer v6 to derive cortical segmentations of 34 ROIs per hemisphere, based on the Desikan-Killiany parcellation scheme^14^. For each T1-MPRAGE data, the pial and white matter surfaces were generated, and the cortical thickness was measured as the distance between the boundaries of pial and white matter surfaces. Visual inspection was carried while blinded to group diagnosis and corrections were performed to ensure accurate skull-stripping and reconstruction of white matter and pial surfaces. Subsequently, PetSurfer was used to segment additional regions, such as the cerebral CSF, pons, skull and air cavities. These segmentations were merged with the default Freesurfer segmentations to facilitate PET-MRI integration and partial-volume correction as previously described^15^. Using the grey matter cerebellum as the reference region, kinetic modelling was performed using the two-stage Multilinear Reference Tissue Model (MRTM2)^16^ within the Petsurfer pipeline to derive partial-volume corrected non-displaceable binding potential (BPND) values for each ROI^15^.

For each subject, we derived a composite PK uptake (global) by averaging the PK binding across each cortical ROI and hemisphere, as well as lobar PK uptake (i.e., frontal, temporal, parietal, occipital). [^11^C]-PiB data were quantified using SUVR with the superior cerebellar grey matter as the reference region. The [^11^C]-PiB SUVR data were similarly subjected to the GTM technique for partial volume correction and treated as a dichotomous variable for Aβ classification (positive if mean SUVR >1.5).

### Diffusion tensor imaging

The DTI acquisition protocol was as follows: 63 slices of 2.0mm thickness, TE = 106 ms, TR = 11700 ms, SENSE = 2, field of view = 192 × 192 mm^2^. The data were preprocessed with the FSL 5.0 software package (http://www.fmrib.ox.ac.uk/fsl). This included registration of all diffusion-weighted images to the b=0 (i.e. no diffusion) volume using the FSL Diffusion Toolbox (*FDT*), followed by brain masks creation with *Brain Extraction Tool* (*BET*), head movement and eddy currents correction. We then used *DTIfit* to independently fit the diffusion tensor for each voxel, resulting in the derivation of FA, MD and RD.

### Statistical analyses

*Tract-based spatial statistics* (*TBSS*) was implemented in the FSL workspace to align each subject’s FA image to a pre-identified target FA image (FMRIB_58)^17^. All aligned FA images were then affine-registered into the Montreal Neurological Institute (MNI) MNI152 template. The mean FA and skeleton were created for all subjects and each FA image was then projected onto the skeleton. The skeleton was thresholded at FA value of 0.2 to include white matter tracts that are common across all subjects, as well as to further exclude voxels that may contain grey matter or CSF. The aligned DTI parameter map of each subject was then projected onto the mean skeleton. In addition, other DTI parameters were aligned by applying the original FA non-linear spatial transformations to the corresponding datasets and projecting them onto the mean FA skeleton. Subsequently, the *randomise* function in FSL was implemented to identify group differences in DTI parameters between DLB and age-matched healthy controls, using a two-sample t-test while controlling for age and gender.

Voxel-wise correlation between DTI and PK11195 was performed with TBSS using age, gender, scan interval between MRI and PET imaging, as well as ACER score as nuisance covariates in the design matrix. Global and lobar PK11195 uptake was treated as the independent variable for each DTI parameter across the white matter skeleton. The statistical significance from these tests was determined using non-parametric permutation testing (n = 5000 permutations)^18^ and adjusted for multiple comparisons by applying the recommended threshold-free cluster enhancement (TFCE p value < 0.05)^19^. Anatomical definitions of significant TBSS clusters were facilitated using the John Hopkins – ICBM white matter atlas, available as part of the FSL package. Finally, the statistical maps were dilated with tbss_fill function for visualisation purposes.

## RESULTS

### Demographics

The demographics and clinical characteristics are reported in Table 1. Both groups were comparable in terms of age, gender and years of education. As expected, MMSE and ACER scores were significantly lower in the DLB group relative to the healthy controls (p < 0.001).

**Table 1.** Clinical characteristics of study sample. Abbreviations: M/F = Male/Female, MMSE = Mini-Mental State Examination, ACE-R = Addenbrooke’s Cognitive Evaluation Revised, * = t-test, # = Mann-Whitney, $ = Chi-Square test.

|  | DLB (n=19) | HC (n=20) | pval |
| --- | --- | --- | --- |
| Age (years) | 73.5±6.1<br>(62-82) | 71.0±6.9<br>(56-81) | 0.23 * |
| Male/Female Ratio | 3.8 (15/4) | 1 (10/10) | 0.06 \$ |
| Education (years) | 12.0±2.3<br>(8-17) | 13.4±2.8<br>(9-19) | 0.10 * |
| MMSE | 22.5±4.3<br>(15-28) | 28.8±1.2<br>(26-30) | <0.0001 # |
| ACE-R | 66.8±11.9<br>(45-87) | 90.9±6.6<br>(75-100) | <0.0001 # |

### Group comparisons of DTI parameters

Voxel-based TBSS analyses revealed that the DLB group relative to controls had significant white matter changes in the body and splenium of corpus callosum. These findings were consistently observed for all three DTI metrics (decreased FA, increased MD and increased RD). In addition, we observed reduced FA and increased RD for DLB in right anterior and posterior corona radiata, left cingulate gyrus, and right superior longitudinal fasciculus (Figure 1). RD alone was increased in the genu of corpus callosum, right internal and external capsule and right uncinate fasciculus. FA was also decreased in bilateral superior part of corona radiata and right posterior thalamic radiations.

**Figure 1:**
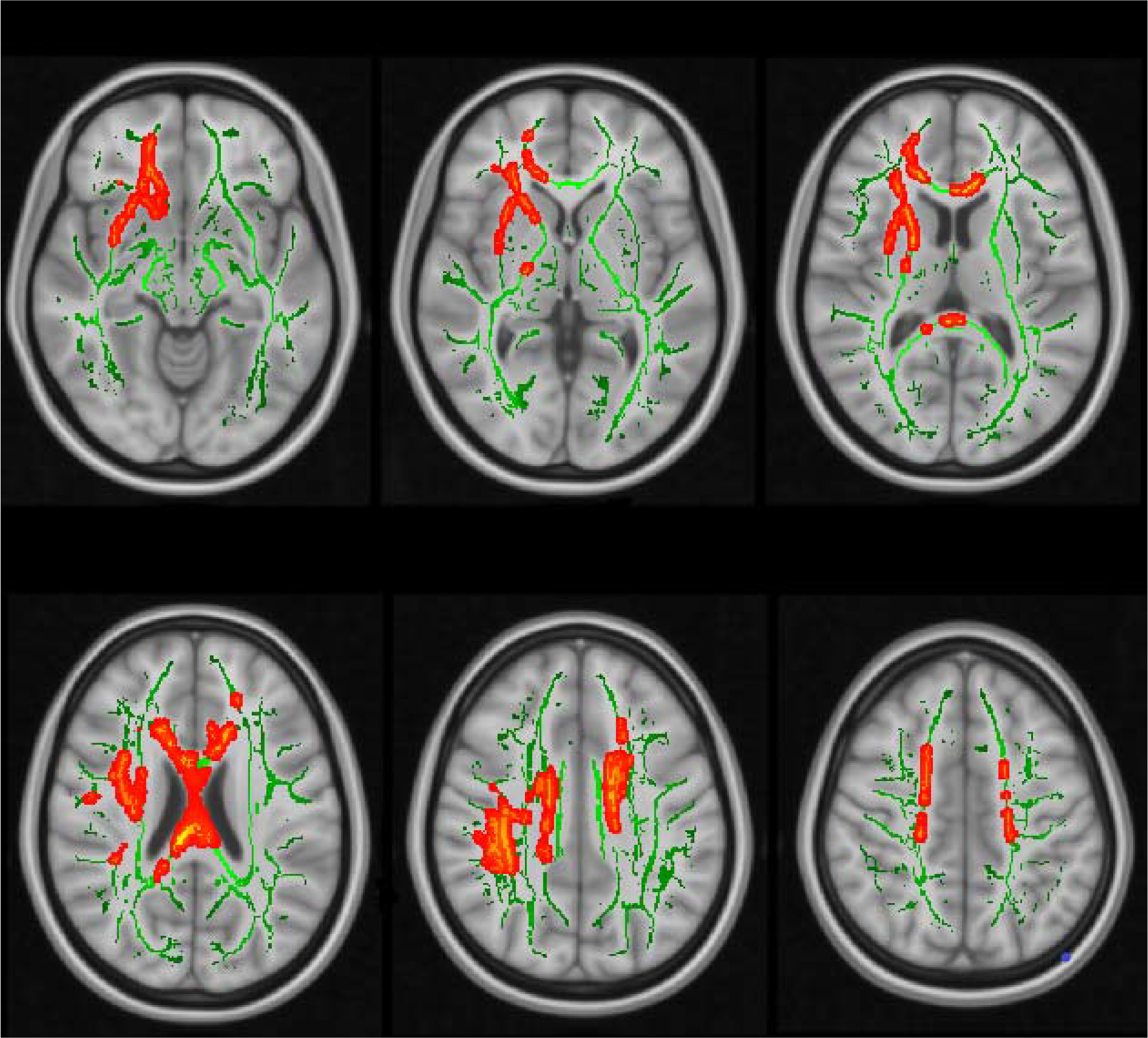
DTI group comparisons showing increased radial diffusivity (red) in DLB subjects compared to healthy control group (TFCE p<0.05)

Subgroup analyses of DLB subjects according to their PiB status revealed a more extensive impairment in the PiB+ DLB (n=9) compared to controls, with virtually every subcortical tract being impaired. In contrast, there was no significant DTI changes in the PiB- DLB subgroup (n=6) compared to controls. Correlation analyses between lobar or global PiB binding and DTI metrics did not reach statistical significance (all p > 0.29). In addition, DTI group comparisons between PiB+ (n = 9) and PiB-DLB (n = 6) did not reach statistical significance (all p > 0.17).

### Voxel-wise association of imaging data

TBSS analyses revealed that higher parietal PK11195 uptake was associated with a relative preservation of white matter (i.e., lower MD and RD) in the corpus callosum (genu and splenium, and to a lesser extent body of corpus callosum), bilateral internal capsule and corona radiata, bilateral posterior thalamic radiations, external capsule and cingulate gyrus, superior longitudinal fasciculus and superior frontooccipital fasciculus, and left sagittal stratum (TFCE p < 0.05) (Figure 2). We did not find significant associations between PK and FA (p = 0.05), nor were there any significant correlations in the opposite contrasts for MD and RD.

**Figure 2:**
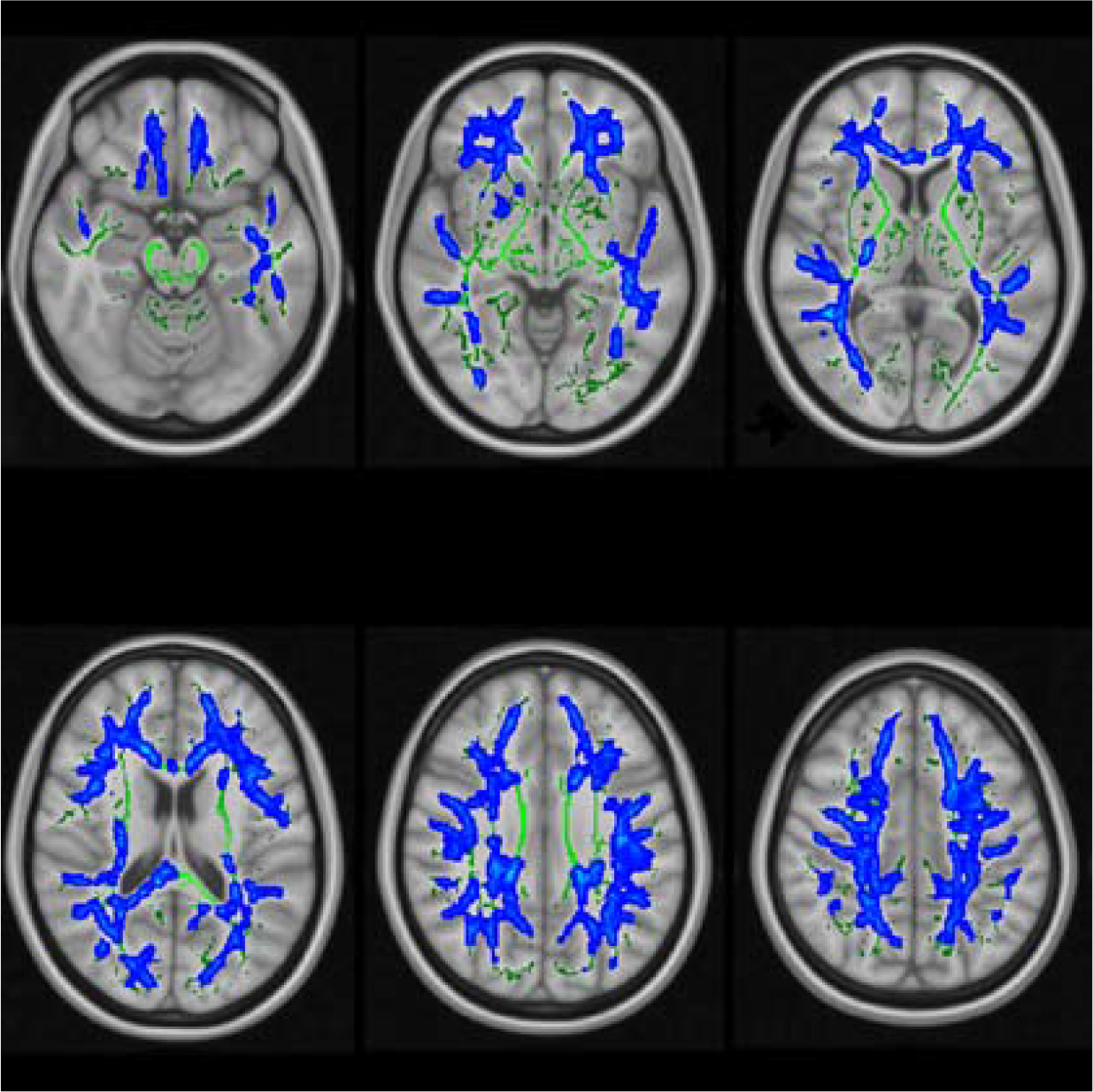
TBSS correlation analyses showing decreased radial diffusivity (blue) correlated with higher parietal ^11^C-PK11195 binding in DLB subjects (TFCE p<0.05)

The analyses did not reach a significant threshold with regards to a correlation between preserved DTI parameters and increased PK11195 at the global (all p > 0.13), frontal (all p > 0.13), temporal (all p > 0.08) and occipital level (all p > 0.13). In addition, higher ACER score was significantly correlated with higher PK11195 binding in frontal (Spearman ρ = 0.52, p = 0.03), temporal (ρ = 0.50, p = 0.03) and occipital lobe (ρ = 0.55, p = 0.02), with a trend for global PK (ρ = 0.46, p = 0.05) but not with parietal PK binding (ρ = 0.26, p = 0.23). There was no significant correlation between lobar and global PiB uptake and DTI parameters (all p > 0.3).

## DISCUSSION

In the present study, we found evidence of relative preservation of white matter integrity in relation to increased microglial activation as measured with PK11195 in DLB. This is consistent with our previous data showing that neuroinflammation in fronto-temporo-parietal cortices was more prominent in DLB subjects with milder cognitive impairment, suggesting that central inflammation plays a pivotal role in the early stages of the disease^6^.

Extensive white matter abnormalities were observed in DLB compared to healthy controls, encompassing the corpus callosum, corona radiata, superior longitudinal fasciculus and white matter underneath left cingulate gyrus, which is in keeping with previous studies^20–24^.

In addition, we observed that higher parietal PK11195 binding was correlated with relative microstructural preservation of white matter tracts (lower diffusivity) in the corpus callosum, internal capsule and corona radiata, superior longitudinal fasciculus and cingulate gyrus, i.e., the same tracts which are impaired in DLB compared to controls. Thus, the spatial congruence of both statistical maps – subjected to the same rigorous statistical thresholds – confer support for the view that white matter deficits may be a secondary event to various processes such as protein deposition, grey matter atrophy and possibly neuroinflammation. In addition, we observed a broad overlap between the findings of MD and RD parameters regarding the correlation of DTI values and PK11195 binding. Higher MD values indicate increased diffusion, suggesting tissue breakdown and increased brain water content. Conversely, higher RD is considered as a more specific marker of myelin damage^25, 26^. As previously reported, MD (and RD) appears as a more sensitive proxy of white matter damage than FA in patients with DLB^23, 24, 27^.

The present study is not without limitations. First, our results have been obtained on a relatively modest sample of subjects with DLB, so this would require confirmation in larger samples. Second, the present relationship between microglial activation and white matter integrity is based on data obtained with a cross-sectional design. Therefore, longitudinal data is required to fully assess the spatial and temporal interplay between white matter integrity and neuroinflammation in DLB. Third, if on the one hand, white matter degeneration can be considered as an early structural marker in neurodegenerative disorders, it can be equally possible that neuroinflammation precipitates white matter damage and thus represents a relatively “upstream” event in the cascade leading to cellular loss in DLB. Consistent with this latter hypothesis, increased microglial activation has been reported in mild cognitive impairment (MCI) and in subjects with REM-sleep behaviour disorder without any motor or cognitive involvement, i.e. conditions preceding the onset of dementia and alpha-synucleinopathies, respectively ^28, 29^. In addition, Femminella *et al.*^30^ showed that increased microglial activation as measured with [^11^C]-PBR28 PET in MCI subjects was associated with higher grey matter and hippocampal volume, suggesting that neuroinflammation is again involved in early stages of the disease or may even play a protective role in the disease trajectory.

The use of PK11195 targeting translocator protein (TSPO) cannot fully represent the extent of central inflammation which is determined by several other factors over and above microglial activation. In fact, other targets such as astrocyte activation should be assessed as they also may play a role in the development of neurodegeneration^31^. In addition, PK11195, as a first generation TSPO ligand, has shown lower sensitivity compared to second generation tracers. However, PK11195 has the advantage of not being affected by the genetic polymorphism (notably rs6971 single nucleotide) altering the binding of second-generation ligands^32, 33^.

In conclusion, we present novel evidence of the *in vivo* association between white matter integrity and microglial activation in DLB. Future longitudinal studies are required to determine whether the early presence of neuroinflammation in DLB recedes and mediates the structural brain abnormalities that we have observed.

## ACKNOWLEDGEMENTS

We are grateful to our volunteers for their participation in the NIMROD study. We thank the radiographers at Wolfson Brain Imaging Centre and the PET/CT unit, Addenbrooke's Hospital for their technical expertise and support in data acquisition. We thank the NIHR Dementias and Neurodegenerative Research Network for their help with subject recruitment. We also thank Dr Istvan Boros, Dr. Joong-Hyun Chun, and WBIC RPU for the manufacture of the [^11^C]-PK11195.

## ETHICS APPROVAL

NRES Committee East of England, Central Cambridge

## REFERENCES

1. Vann Jones SA, O'Brien JT. The prevalence and incidence of dementia with Lewy bodies: a systematic review of population and clinical studies. Psychol Med 2014;44(4):673–683.

2. Spillantini MG, Schmidt ML, Lee VM, Trojanowski JQ, Jakes R, Goedert M. Alpha-synuclein in Lewy bodies. Nature 1997;388(6645):839–840.

3. Gomperts SN. Lewy Body Dementias: Dementia With Lewy Bodies and Parkinson Disease Dementia. Continuum (Minneap Minn) 2016;22(2 Dementia):435–463.

4. Turkheimer FE, Edison P, Pavese N, et al. Reference and target region modeling of [11C]-(R)-PK11195 brain studies. J Nucl Med 2007;48(1):158–167.

5. Iannaccone S, Cerami C, Alessio M, et al. In vivo microglia activation in very early dementia with Lewy bodies, comparison with Parkinson's disease. Parkinsonism Relat Disord 2013;19(1):47–52.

6. Surendranathan A, Su L, Mak E, et al. Early microglial activation and peripheral inflammation in dementia with Lewy bodies. Brain 2018;141(12):3415–3427.

7. Hamelin L, Lagarde J, Dorothee G, et al. Early and protective microglial activation in Alzheimer's disease: a prospective study using 18F-DPA-714 PET imaging. Brain 2016;139(Pt 4):1252–1264.

8. Passamonti L, Rodriguez PV, Hong YT, et al. [(11)C]PK11195 binding in Alzheimer disease and progressive supranuclear palsy. Neurology 2018;90(22):e1989–e1996.

9. Fan Z, Okello AA, Brooks DJ, Edison P. Longitudinal influence of microglial activation and amyloid on neuronal function in Alzheimer's disease. Brain 2015;138(Pt 12):3685–3698.

10. Bevan-Jones WR, Surendranathan A, Passamonti L, et al. Neuroimaging of Inflammation in Memory and Related Other Disorders (NIMROD) study protocol: a deep phenotyping cohort study of the role of brain inflammation in dementia, depression and other neurological illnesses. BMJ Open 2017;7(1):e013187.

11. McKeith IG, Dickson DW, Lowe J, et al. Diagnosis and management of dementia with Lewy bodies: third report of the DLB Consortium. Neurology 2005;65(12):1863–1872.

12. McKeith IG, Boeve BF, Dickson DW, et al. Diagnosis and management of dementia with Lewy bodies: Fourth consensus report of the DLB Consortium. Neurology 2017;89(1):88–100.

13. Jagust WJ, Bandy D, Chen K, et al. The Alzheimer's Disease Neuroimaging Initiative positron emission tomography core. Alzheimers Dement 2010;6(3):221–229.

14. Desikan RS, Segonne F, Fischl B, et al. An automated labeling system for subdividing the human cerebral cortex on MRI scans into gyral based regions of interest. Neuroimage 2006;31(3):968–980.

15. Greve DN, Svarer C, Fisher PM, et al. Cortical surface-based analysis reduces bias and variance in kinetic modeling of brain PET data. Neuroimage 2014;92:225–236.

16. Ichise M, Liow JS, Lu JQ, et al. Linearized reference tissue parametric imaging methods: application to [11C]DASB positron emission tomography studies of the serotonin transporter in human brain. J Cereb Blood Flow Metab 2003;23(9):1096–1112.

17. Smith SM, Jenkinson M, Woolrich MW, et al. Advances in functional and structural MR image analysis and implementation as FSL. Neuroimage 2004;23 Suppl 1:S208–219.

18. Nichols TE, Holmes AP. Nonparametric permutation tests for functional neuroimaging: a primer with examples. Hum Brain Mapp 2002;15(1):1–25.

19. Smith SM, Nichols TE. Threshold-free cluster enhancement: addressing problems of smoothing, threshold dependence and localisation in cluster inference. Neuroimage 2009;44(1):83–98.

20. Firbank MJ, Blamire AM, Krishnan MS, et al. Diffusion tensor imaging in dementia with Lewy bodies and Alzheimer's disease. Psychiatry Res 2007;155(2):135–145.

21. Firbank MJ, Watson R, Mak E, et al. Longitudinal diffusion tensor imaging in dementia with Lewy bodies and Alzheimer's disease. Parkinsonism Relat Disord 2016;24:76–80.

22. Kantarci K, Avula R, Senjem ML, et al. Dementia with Lewy bodies and Alzheimer disease: neurodegenerative patterns characterized by DTI. Neurology 2010;74(22):1814–1821.

23. Delli Pizzi S, Franciotti R, Taylor JP, et al. Structural Connectivity is Differently Altered in Dementia with Lewy Body and Alzheimer's Disease. Front Aging Neurosci 2015;7:208.

24. Watson R, Blamire AM, Colloby SJ, et al. Characterizing dementia with Lewy bodies by means of diffusion tensor imaging. Neurology 2012;79(9):906–914.

25. Pierpaoli C, Basser PJ. Toward a quantitative assessment of diffusion anisotropy. Magn Reson Med 1996;36(6):893–906.

26. Song SK, Yoshino J, Le TQ, et al. Demyelination increases radial diffusivity in corpus callosum of mouse brain. Neuroimage 2005;26(1):132–140.

27. O'Donovan J, Watson R, Colloby SJ, Blamire AM, O'Brien JT. Assessment of regional MR diffusion changes in dementia with Lewy bodies and Alzheimer's disease. Int Psychogeriatr 2014;26(4):627–635.

28. Okello A, Edison P, Archer HA, et al. Microglial activation and amyloid deposition in mild cognitive impairment: a PET study. Neurology 2009;72(1):56–62.

29. Stokholm MG, Iranzo A, Ostergaard K, et al. Assessment of neuroinflammation in patients with idiopathic rapid-eye-movement sleep behaviour disorder: a case-control study. Lancet Neurol 2017;16(10):789–796.

30. Femminella GD, Dani M, Wood M, et al. Microglial activation in early Alzheimer trajectory is associated with higher gray matter volume. Neurology 2019.

31. Santillo AF, Gambini JP, Lannfelt L, et al. In vivo imaging of astrocytosis in Alzheimer's disease: an (1)(1)C-L-deuteriodeprenyl and PIB PET study. Eur J Nucl Med Mol Imaging 2011;38(12):2202–2208.

32. Owen DR, Yeo AJ, Gunn RN, et al. An 18-kDa translocator protein (TSPO) polymorphism explains differences in binding affinity of the PET radioligand PBR28. J Cereb Blood Flow Metab 2012;32(1):1–5.

33. Kreisl WC, Jenko KJ, Hines CS, et al. A genetic polymorphism for translocator protein 18 kDa affects both in vitro and in vivo radioligand binding in human brain to this putative biomarker of neuroinflammation. J Cereb Blood Flow Metab 2013;33(1):53–58.

